# A generalizable protocol for expression and purification of membrane-bound bacterial phosphoglycosyl transferases in liponanoparticles

**DOI:** 10.1101/2023.03.20.533523

**Authors:** Greg J. Dodge, Hannah M. Bernstein, Barbara Imperiali

**Author notes:** Equal contributions (Greg J. Dodge, Hannah M. Bernstein).

## Abstract

Phosphoglycosyl transferases (PGTs) are among the first membrane-bound enzymes involved in the biosynthesis of bacterial glycoconjugates. Robust expression and purification protocols for an abundant subfamily of PGTs remains lacking. Recent advancements in detergent-free methods for membrane protein solubilization open the door for purification of difficult membrane proteins directly from cell membranes into native-like liponanoparticles. By leveraging autoinduction, *in vivo* SUMO tag cleavage, styrene maleic acid co-polymer liponanoparticles (SMALPs), and Strep-Tag purification, we have established a robust workflow for expression and purification of previously unobtainable PGTs. The material generated from this workflow is extremely pure and can be directly visualized by Cryogenic Electron Microscopy (CryoEM). The methods presented here promise to be generalizable to additional membrane proteins recombinantly expressed in *E. coli* and should be of interest to the greater membrane proteomics community.

**Highlights:**

- Expression and purification of full-length Lg-PGTs has proven challenging.
- Autoinduction and *in vivo* Ulp1 cleavage produces active full-length Lg-PGTs.
- SMA and DIBMA are vastly superior to DDM for Lg-PGT solubilization.
- Strep-tag purification yields SMALPs suitable for CryoEM characterization.

## Introduction

The presence of a heterogenous coat of cell surface and extracellular glycoconjugates is a common theme across all domains of life.[1] Notable examples of glycoconjugates in prokaryotes include lipopolysaccharide (LPS), capsular polysaccharide (CPS), exopolysaccharide (EPS), and wall teichoic acid (WTA). Biosynthesis of glycoconjugates in prokaryotes can be broadly classified into the wzx/wzy-dependent or wzx/wzxy-independent pathways.[2, 3] In bacteria, including many human pathogens, genetic deletion or small molecule-based inhibition of the biosynthetic machinery involved in the assembly of the complex glycoconjugates reduces viability and impacts virulence.[4, 5] As such, the enzymes involved in glycoconjugate biosynthesis represent attractive targets for detailed studies.

Among the enzymes involved in the early steps of *en bloc* glycoconjugate biosynthesis in bacteria are phosphoglycosyl transferases (PGTs).[6, 7] These enzymes catalyze the transfer of a phospho-sugar from a nucleotide sugar donor to an acceptor polyprenol phosphate – most commonly undecaprenol phosphate (UndP).[8] PGTs are integral membrane proteins and can be classified into two superfamilies based on the topologies of the functional domains – polytopic (polyPGTs) wherein the catalytic domain comprises several transmembrane helices (TMH) and monotopic (monoPGTs), with a catalytic domain including a single reentrant membrane helix (RMH).[6, 9] Initial sequence and hydropathy analysis of monoPGTs misannotated the RMH as a TMH, confounding early topological assessment of these enzymes.[10, 11] Landmark studies culminating with the crystal structure of PglC from *Campylobacter concisus*, unequivocally established the topology of the monoPGT catalytic domain and identified a distinct sequence fingerprint for the RMH and catalytic motif.[12–14]

Recently, the PGT sequence fingerprint was utilized to establish a sequence similarity network (SSN) comprising 38,000 nonredundant PGT sequences.[15] Analysis of this network demonstrated that monotopic PGTs broadly cluster into three sub-families: small (Sm-PGT), containing only the catalytic domain, bifunctional (Bi-PGT) containing the catalytic PGT domain and an additional functional domain, and large (Lg-PGT) containing a putative four-transmembrane helix bundle and a domain of unknown function (DUF) fused to the N-terminus of the catalytic domain. A comparison of the domain structure of the Sm- and Lg-PGTs is illustrated in Figure 1. The Lg-PGT subfamily is predominant and represents 47% of all non-redundant PGT sequences within the SSN. However, biochemical characterization of these proteins remains limited as recombinant expression of the full-length integral membrane Lg-PGTs has represented a major hurdle to progress in understanding the extensive enzyme family. Additionally, the establishment of the PGT catalytic domain topology warrants careful re-examination of early studies, as the proper orientation of Lg-PGTs places the DUF found N-terminal to the catalytic domain in the cytoplasm as opposed to the periplasm.[10, 11, 14, 16]

**Figure 1.**
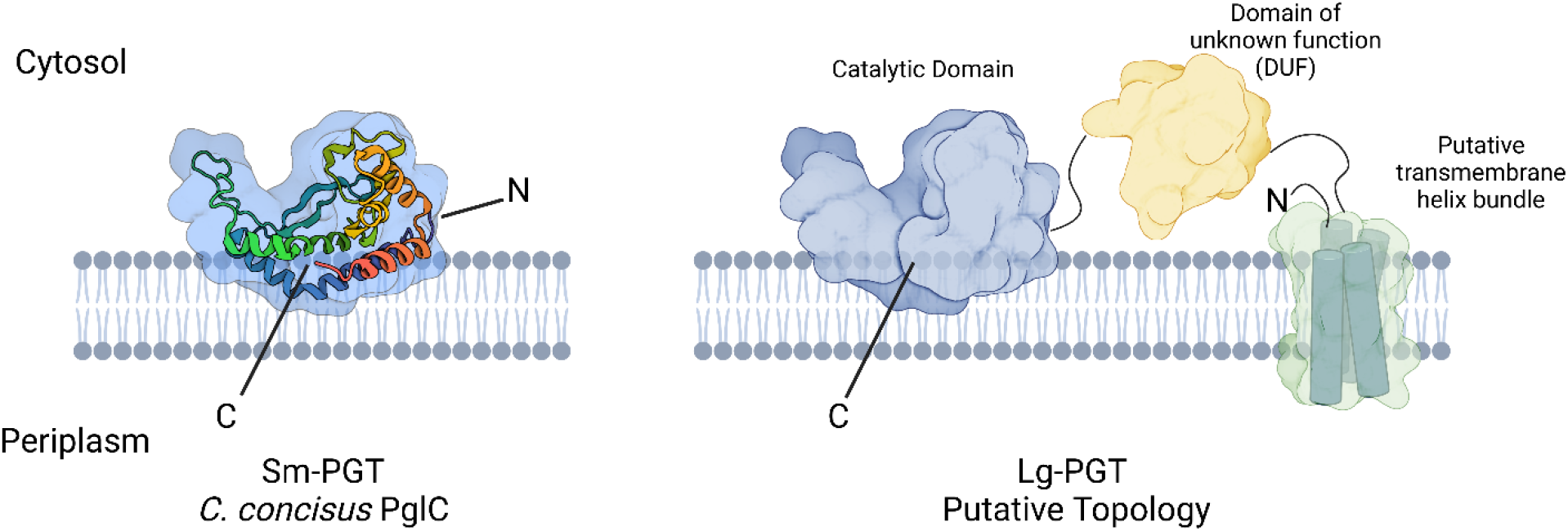
Comparison of topology between the Sm- and Lg-PGTs. Left: PglC from *Campylobacter concisus* (PDBID: 5W7L) comprises only the PGT catalytic domain, enters and exits the membrane on the same leaflet, by way of the N-terminal re-entrant membrane helix (RMH). Right: Schematic of a LG-PGT. The N-terminus of Lg-PGT contains four putative transmembrane helices, followed by a domain of unknown function, and finally a catalytic domain resembling PglC.

Methodologies for stabilizing membrane proteins on extraction from cell membrane are rapidly evolving.[17] Historically, detergents such as n-dodecyl-β-D-maltoside (DDM) or N,N-dimethyl-n-dodecylamine N-oxide (LDAO) have been used to solubilize membrane-bound proteins into detergent micelles, often at the expense of protein stability.[18] Recently, techniques for isolating target proteins within liponanoparticles have been described. Of these, the styrene maleic acid copolymer (SMA), stands out for its ability to extract proteins directly into liponanoparticles known as SMALPs.[19] SMALPs are compatible with common purification techniques,[20, 21] often stabilize the targets they encapsulate,[22] and have been used as a platform for a wide variety of studies.[23–26]

Here, we describe an optimized expression system, for recombinant production of full-length Lg-PGTs from various bacteria (Fig. 2). Using radioactivity-based assays, we investigate the substrate specificity of successfully expressed targets in cell envelope fractions. Then, we explore the application of the SMALP approach for solubilizing Lg-PGTs and identify optimal conditions for producing highly purified liponanoparticles which can be directly visualized via cryogenic electron microscopy (CryoEM). These studies establish SMALP as powerful approach for Lg-PGT purification, highlight the substrate specificity of recombinantly expressed Lg-PGTs, and establish the groundwork for future biophysical and structural biology studies of this major subfamily of monoPGTs. Significantly, the overall strategy may prove advantageous in applications towards other classes of integral membrane proteins.

**Figure 2.**
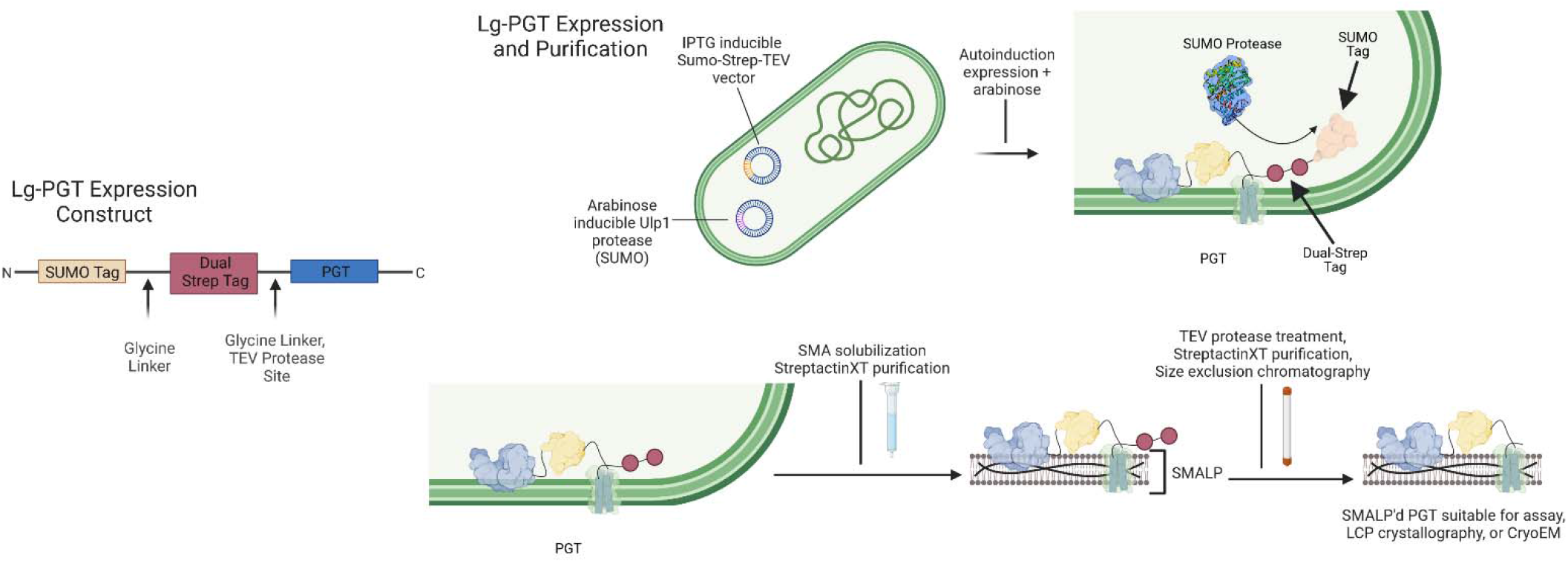
Top: Lg-PGT expression and purification strategy. Left: The expression vector encodes an N-terminal SUMO fusion, a dual-strep tag for purification, a TEV cleavage site for tag removal, followed by Lg-PGTs of interest. Right: The expression and purification strategy utilizes co-expression of the Ulp1 protease for *in vivo* SUMO tag removal. Downstream chromatography utilizing StreptactinXT resin provides highly pure material. Optional clean-up can be achieved using size exclusion chromatography, and the dual-strep tag can be cleaved via incubation with TEV protease.

## Materials and Methods

Synthetic genes encoding WbaP from S. *enterica* (Uniprot: P26406), WbaP from *T. thermophilus* (Uniprot: A0A510HWX9), WecP from *Aeromonas hydrophila* (Uniprot: B3FN88), CpsE from *Streptococcus pneumoniae* (Uniprot: Q8KW9P) and WcaJ from *Escherichia coli* (Uniprot: P71241) were codon optimized for expression in *E. coli*. An expression vector was derived from pE-SUMOpro (LifeSensors) to encode a dual-strep tag followed by a TEV protease site between the SUMO tag and MCS (His-SUMO-dual strep-TEV-PGT). Fragments were assembled using Gibson assembly. All plasmid sequences were verified via sanger sequencing. Plasmids were transformed into *E. coli* C43 harboring pAM174,[27] which encodes Ulp1 under an arabinose promotor. Proteins were expressed using autoinduction[28] in 0.5 L cultures supplemented with 150 μg/mL kanamycin and 25 μg/mL chloramphenicol at 37°C. One gram of powdered L-arabinose was added to cultures after OD_600_ reached ~1.5 and the temperature was adjusted to 18° C. After 18 hr expression, cells were harvested via centrifugation, transferred to 1-gallon Ziplock^®^ freezer bags, spread to a thin layer, and frozen at −80° C.

To express SUMO-tagged CpsE and WcaJ, the above protocol was followed using BL21 DE3 *E. coli* lacking the pAM174 plasmid.

### Protein Purification

#### Isolation of Cell membranes

All purification steps were performed on ice unless otherwise noted. Frozen cell pellets were manually broken in bags and resuspended in buffer A (50 mM HEPES pH 8.0, 300 mM NaCl) at 4 mL per g pellet. Resuspended pellets were supplemented with 2 mM MgCl_2_, 0.06 mg/mL lysozyme (RPI), and 0.5 mg/mL DNase I (Millipore Sigma) and incubated for 30 minutes. Cells were disrupted via sonication (2x, 50% amplitude, 1s on 2 s off), and cell membrane was isolated via differential centrifugation (Ti-45 rotor, 9,000g 45 min, reserve supernatant, 140,000g 65 min). Cell membranes were resuspended to 50 mg/mL (assessed by raw UV signal at 280 nm) in buffer A, a 1 mL aliquot was reserved for activity assay, flash frozen in liquid N_2_, and stored at −80° C.

#### Catalytic variants of E. coli WcaJ and S. pneumoniae CpsE

Site-directed mutagenesis was done via PCR. Primers were designed via Quikchange primer design tool (Agilent). Designed primer pairs are shown below. Sequences of all mutagenized plasmids were confirmed by Sanger sequencing.

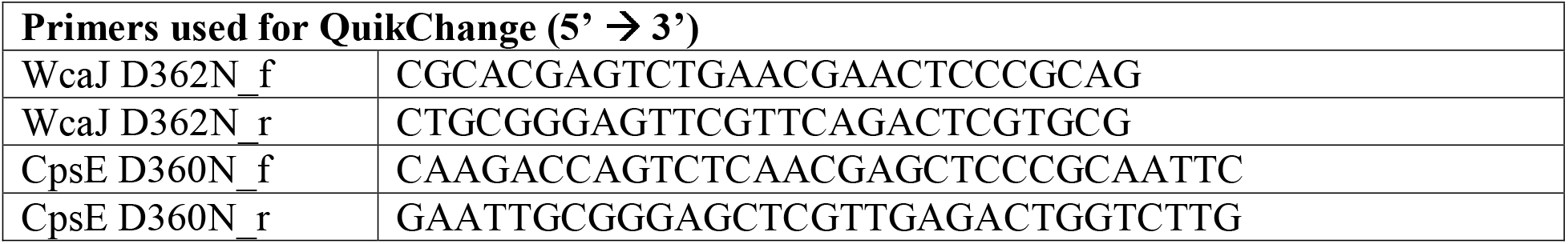

#### SMA solubilization and purification

The frozen cell membrane fraction was thawed and mixed 1:1 (v/v) with a 2% stock solution of styrene maleic acid copolymer (Polyscope) or diisobutylene maleic acid copolymer (BASF) in buffer A and rotated at room temperature for one hour. Soluble SMA liponanoparticles (SMALPs) or DIBMA liponanoparticles (DIBMALPs) were isolated by centrifugation (Ti45 rotor, 160,000g 65 min). The supernatant was flowed over 1 mL StreptactinXT 4flow resin (IBA Biosciences) pre-equilibrated with buffer A. The flow-through was re-run over the column bed, and the column was washed with 5 CV buffer A. Protein was eluted using 3 CV Buffer A + 50 mM Biotin (buffer B). Protein containing fractions were pooled and loaded onto 3x tandem 5 mL HiTrap desalting columns (Cytiva) equilibrated with 25 mM HEPES pH 8.0, 150 mM NaCl (buffer C) to remove biotin. Protein purity was assessed via SDS-PAGE. For SMALP samples, protein concentration was determined via BCA assay (Pierce). Protein was concentrated to 2 mg/mL, and flash frozen in liquid N_2_, then stored at −80° C.

### Activity assay

The PGT radioactivity-based activity assay followed previously published protocols.[29] Assay buffer comprised 20 μM UndP, 0.1% Triton X-100, 10% DMSO, 50 mM HEPES pH 7.5, 100 mM NaCl, 5 mM MgCL_2_, 20 μM UndP, 2 μL [^3^H] labeled UDP-sugar substrate (Table S1). A 4 μL aliquot of PGT-containing CEF was added to 46 μL reaction buffer and incubated at room temperature. A series of 12 μL aliquots were taken at 0.5, 1, 3, 6 min, and quenched by addition to a vial containing 1 mL 2:1 chloroform:methanol and 500 μL pure solvent upper phase (PSUP, 15 mL chloroform, 240 mL methanol, 1.83 g KCl, 235 mL H_2_O). Quenched reactions were vortexed, aqueous/organic phases were allowed to separate, and the aqueous phase was removed. Reactions were re-extracted an additional 2x with 500 μL PSUP. A 5 mL aliquot of Opti-Fluor™ O scintillation fluid (Perkin Elmer) was added to each reaction, and tubes were counted on a liquid scintillation counter (Beckman LS6500). Readings were taken in triplicate, and figures were generated using GraphPad Prism (Version 9.4.1 for Windows, GraphPad Software, San Diego, California USA, www.graphpad.com”).

To measure activity of SUMO-tagged CpsE and WcaJ, the above protocol was modified to include 1 μM non-radiolabeled UDP-Glc in the assay buffer.

#### Cryo-Electron Microscopy of S. enterica WbaP in SMALP

Quantafoil 1.2/1.3 300 mesh Cu grids were glow discharged using an EMITech K100 glow discharger at 15 mA for 60 seconds. Grids were prepared using a Vitrobot Mark IV (Thermo Scientific) following standard protocols. Briefly, 2.5 μL of S. *enterica* WbaP in SMA30 at 2.0 mg/mL was deposited on a glow-discharged grid. After 5 seconds, grids were blotted at blot force 0 for 7 seconds and plunged into liquid ethane. Grids were clipped and stored in liquid N_2_.

Movies were collected on a 200 kV Talos Arctica equipped with a Falcon 3EC camera at 73,000x magnification, nominal pixel size of 2.007 Å, with a spherical aberration of 2.7 mm. Total dose on the sample was 56.75 e/Å^2^. A total of 544 movies were collected, and data were processed using Cryosparc V 3.0.1.[30]

Figures generated using Biorender.com

## Results

### Expression and purification of full-length dual-strep tagged Lg-PGTs in styrene maleic acid liponanoparticles (SMALP)

Initial attempts at the expression and solubilization of full-length Lg-PGTs using DDM yielded only poor yields of truncated protein (Fig S1). Previously, our group had successfully applied detergent-free methods to study Sm-PGTs.[23, 24] Accordingly, we hypothesized that isolating target Lg-PGTs in liponanoparticles, as applied to Sm-PGT could provide an alternative approach for stabilization and purification. Owing to the capability to directly extract membrane proteins from lipids in a bacterial cell envelope fraction, we turned to SMA and related DIBMA polymers.[20, 31] As these polymers have only recently been described, robust protocols to produce material suitable for applications such as structural and functional studies continue to be explored. We developed the expression and purification scheme based on several criteria: 1: Compatibility with a SUMO tag, which promotes improved expression yields and the stability of membrane proteins,[32] 2: Compatibility with autoinduction,[28] which has previously been used to express robust levels of PGTs,[8, 12] 3: Use of a purification strategy that avoids immobilized metal ion affinity chromatography (IMAC), because the SMA and DIBMA polymers interact with divalent cations such as Ni^2+^ hindering purification.[25, 33, 34]

An expression plasmid was constructed that encoded an N-terminal SUMO-tag, a short linker followed by a dual-strep tag, a short linker preceding a tobacco etch virus (TEV) protease cleavage site, and the genes of interest (Figure 2). Additionally, a modified C43-based *E. coli* expression system was chosen which facilitated *in vivo* cleavage of the SUMO tag by co-expression of Ulp1 under an orthogonal promotor (pBAD).[27]

A small panel of SMA polymers was chosen to screen for optimal solubilization of Lg-PGTs. The SMA polymers differ from one another in terms of the syrene:maleic acid ratio, while DIBMA contains a diisobutylene moiety in place of the styrene found in the SMA polymers. Solubilized material was then purified using StreptactinXT resin, and SDS-PAGE was used to assess solubilization and purification efficiency. As is often the case with SMA-solubilization, each of the PGTs tested displayed differing solubilization profiles (Fig. 3). At least one condition for solubilization and purification was identified for each of the proteins assessed, with SMA-30, in general, performing well for all Lg-PGTs.

**Figure 3.**
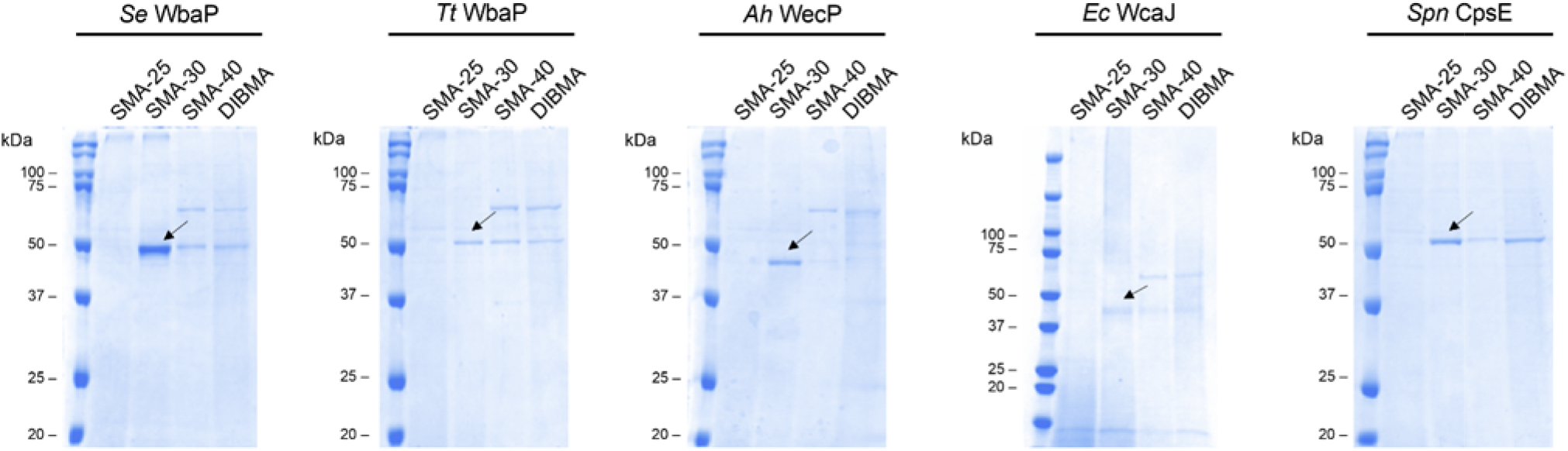
SDS-PAGE gels of elution fractions from small-scale SMALP purification screen. Arrows denote bands corresponding to purified target proteins. A small set of SMA polymers were screened, with SMA30 demonstrating optimal solubilization for each of the targets. Owing to their high isoelectric points, each PGT runs at a lower-than-expected molecular weight. Predicted molecular weights (kDa): *Se* WbaP: 61.2, *Tt* WbaP: 58.7, *Ah* WecP 54.0, *Ec* WcaJ: 57.5, *Spn* CpsE: 57.2.

### Activity and substrate specificity of Lg-PGTs in cell membranes

The ability to heterologously express full-length Lg-PGTs from various organisms in *E. coli* provided enzymes to assess NDP-sugar substrate selectivity. A radioactivity-based assay was used to monitor transfer of labeled phospho-sugars onto UndP.[29] Recombinantly expressed WbaP from S. *enterica* exhibited high specificity for UDP-Gal, as did WbaP from *T. thermophilus* (Fig. 4). WecP from *A. hydrophila* was found to utilize UDP-GalNAc as its preferred substrate (Fig. 4). In initial analyses, both *E. coli* WcaJ and S. *pneumoniae* CpsE expressed in C43 *E. coli* did not show turnover for any of the substrates paneled, despite reports of both of these enzymes utilizing UDP-Glc (Fig. S3).[35, 36]

**Figure 4.**
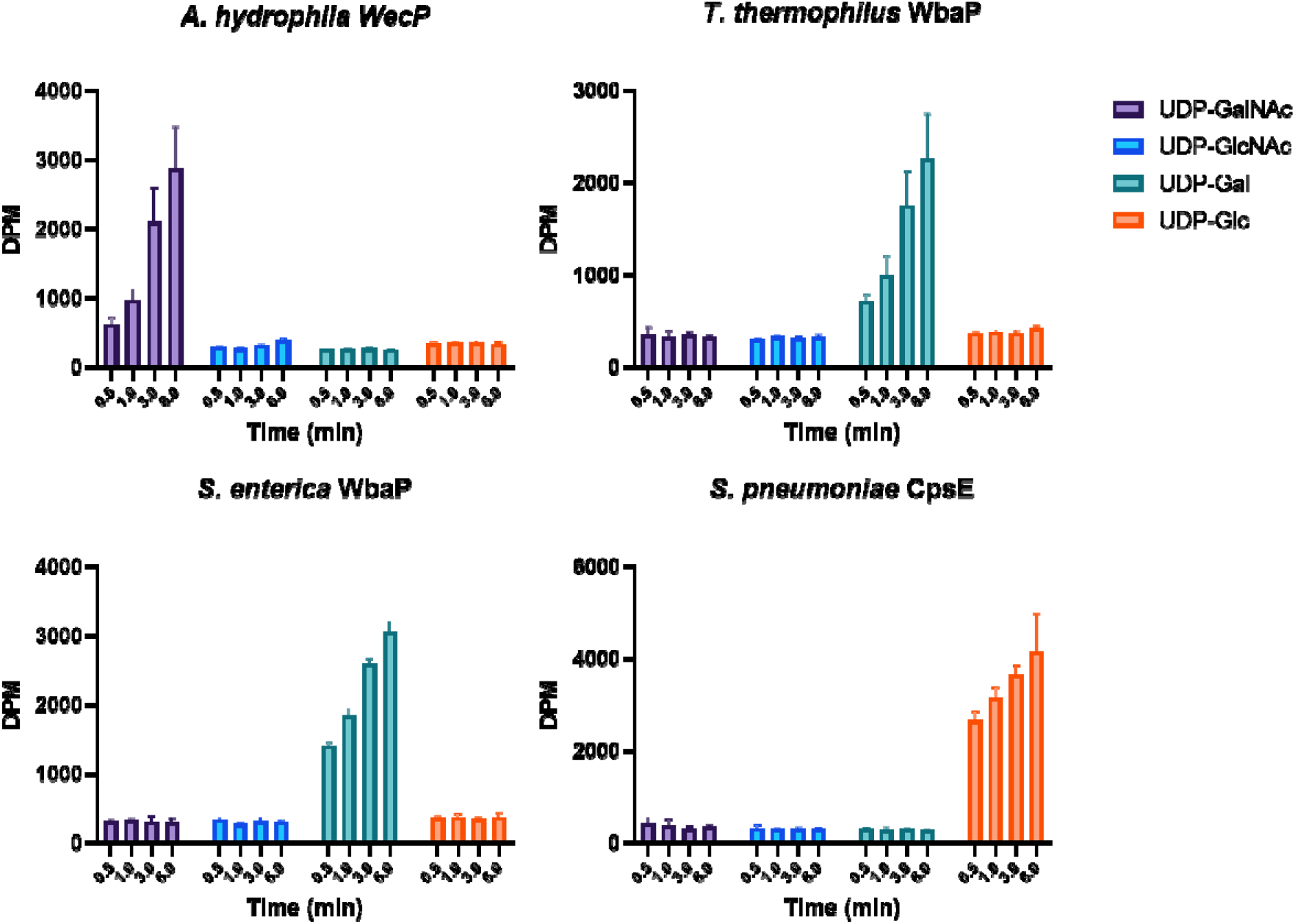
Substrate Panel of full-length LG-PGTs in cell membranes. S. *enterica* and *T. thermophilus* WbaP in cell membranes display high specificity for UDP-Gal. *A. hydrophila* WecP displays high specificity for UDP-GalNAc. S. *pneumoniae* CpsE displays no activity in C43 cell membranes, but readily transfers UDP-Glc when expressed as a SUMO-fusion in BL21 DE3.

In effort to identify conditions in which active WcaJ and CpsE could be produced, we expressed these plasmids in BL21 DE3 *E. coli*, which would leave the potentially stabilizing SUMO-tag in place. This led to the identification of an unusual phenotype. When overexpressed in BL21 DE3 *E. coli*, both SUMO-WcaJ and SUMO-CpsE caused production of a viscous, opaque slime visible after cells were pelleted (Fig. S2). This behavior was not observed when variants lacking the catalytic aspartic acid in the conserved DE dyad were overexpressed.[8, 13] Owing to the striking mucoid phenotype in cell culture, we isolated BL21 DE3 cell membranes containing SUMO-CpsE and SUMO-WcaJ and assessed activity using UDP-Glc as a substrate. Transfer of [^3^H]-labeled UDP-Glc to UndP was observed in both experiments, although WcaJ had reduced activity compared to the other Lg-PGTs assayed (Fig. 4, S3). The ability to monitor activity of WcaJ and CpsE using OD of spent media will greatly enhance future inhibitor development by providing a simple and straightforward *in vivo* activity readout. Retention of the SUMO-tag during expression provides a route to recombinantly express Lg-PGTs that may otherwise prove inactive or unstable.

### Cryo-Electron Microscopy (CryoEM) of Lg-PGTs in SMALP

Although the molecular weights of the Lg-PGTs expressed using the presented Strep-tag/SMALP system are relatively small for CryoEM (~60 kDa), we reasoned that the additional mass from the stabilizing SMA polymer and the phospholipid annulus around the proteins in SMALP may facilitate direct visualization of Lg-PGTs. As S. *enterica* WbaP solubilized in SMA30 gave highest yields, this protein was chosen for CryoEM analysis. Particles were readily visible in micrographs collected, and 2D classes generated from a small dataset showed intriguing features, including putative C2 symmetry (Fig. 5). Efforts to solve a high-resolution reconstruction of S. *enterica* WbaP in SMALP will be reported in due course. These results will yield critical information on the positioning of Lg-PGTs within the lipid bilayer, as well as the overall architecture of these complex multi-domain enzymes.

**Figure 5.**
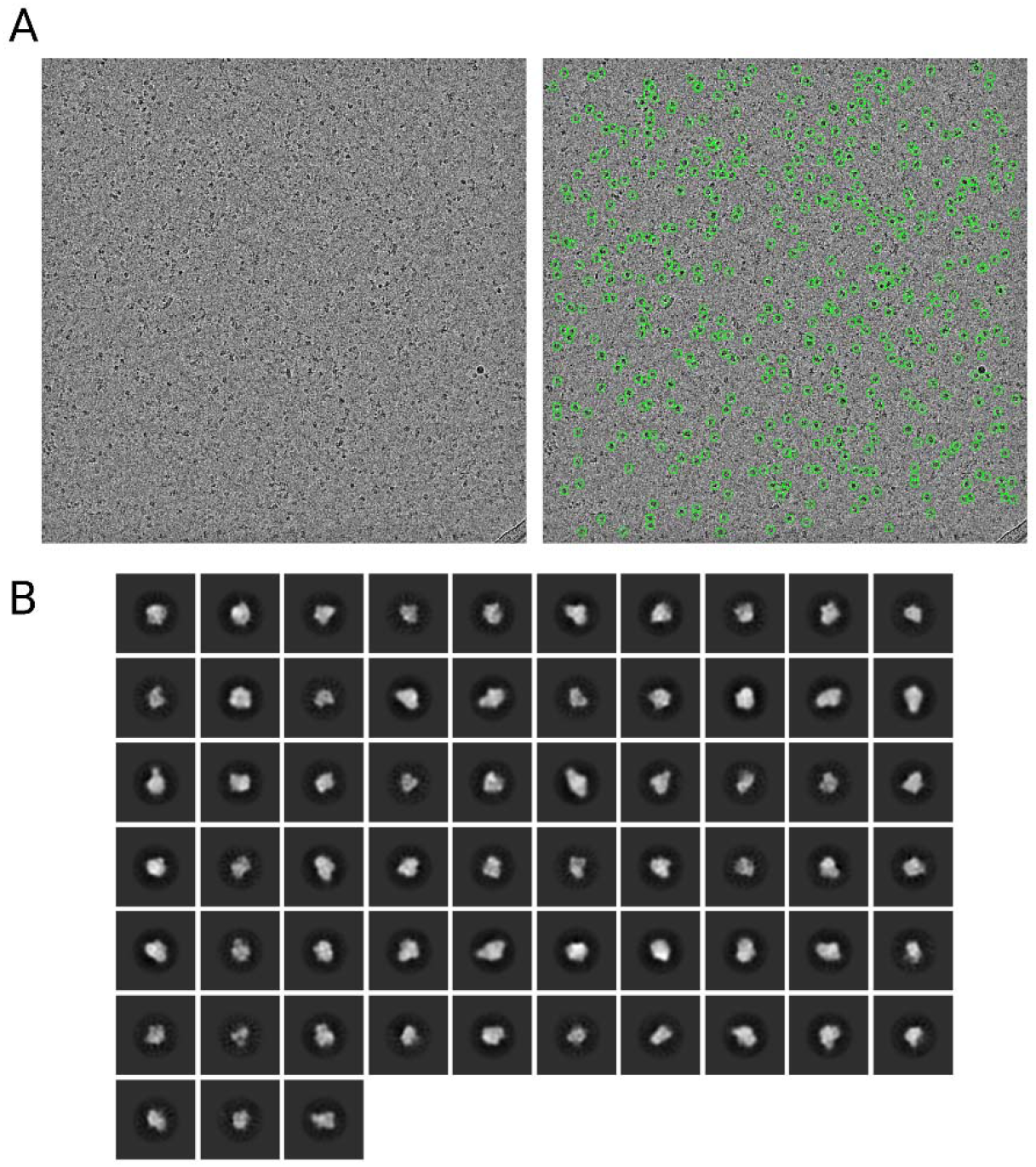
Cryogenic Electron microscopy of purified S. *enterica* WbaP in SMA30 liponanoparticles. Top: Example micrograph lowpass filtered to 5Å and particles picked in cryosparc. Bottom: Selected 2D class averages from 189,401 extracted particles.

## Discussion

Bacterial glycoconjugates play critical roles survival, virulence, antibiotic resistance, and host immunomodulation. However, understanding their biosynthesis is stymied by the non-templated nature of their assembly. Furthermore, linking substrate specificity to sequence of enzymes involved in glycoconjugate assembly has proven a significant challenge. As such, direct characterization of the enzymes involved in glycoconjugate biosynthesis remains a requirement to fully understand pathway logic. Compounding the above issues is the localization of many of the glycoconjugate assembly enzymes to the membrane. Methodologies to improve expression and purification of membrane-bound glycoconjugate biosynthesis enzymes will advance studies connecting enzyme sequence, to structure, function, and final glycoconjugate identity.

Previous attempts to recombinantly overexpress Lg-PGTs has been met with limited success.[35, 37–40] Here we present an improved method of producing full-length Lg-PGTs in C43 *E. coli* by incorporating a SUMO-tag for stabilization and improved yield, utilizing autoinduction, and *in vivo* Ulp1 cleavage to remove the SUMO tag from mature protein prior to extraction of PGTs from cell membranes (Figs. 2, 3). Cell membranes isolated from *E. coli* overexpressing S. *enterica* WbaP, and *A. hydrophila* WecP contained active full-length enzymes whose substrate preference matches that reported previously (Fig. 4). *T. thermophilus* WbaP was also active in cell membranes, and displayed strong preference for UDP-Gal, implying that the sugar at the reducing end of LPS O-Antigen repeat unit for strain ATCC 27634 is Gal. Both *E. coli* WcaJ and S. *pneumoniae* CpsE were inactive in membranes isolated from C43 *E. coli* (Figs. 4, S3). Intriguingly, when expressed in BL21 DE3 lacking co-expressed Ulp1, both plasmids caused the appearance of a turbid mucoid slime in the media. Membranes isolated from these cells readily transferred UDP-Glc onto UndP *in* vitro (Figs. 4, S3). The observed phenotype is reminiscent of cells producing colanic acid.[41] Additionally, when the signature active site aspartic acid residue in either WcaJ or CpsE were mutated to asparagine, the spent media remined optically transparent after cells were removed via centrifugation. As Und-PP-Glc is the initial lipid-linked glycan in colanic acid biosynthesis,[42] it is tempting to speculate that the mucoid phenotype may be a result of colanic acid overproduction in effort to process UndP and Und-PP-Glc through the pathway to recycle the essential UndP.

Although detergent solubilization has been a mainstay for membrane protein extraction and stabilization, the application of detergents comes with significant downsides. Namely, excess detergent above the critical micelle concentration is required in all buffers after protein extraction, detergent micelles often destabilize solubilized proteins in part due to the depletion of the native phopholiplipid context, and some membrane proteins remain recalcitrant to detergent solubilization. Indeed, in the case of each of the Lg-PGTs tested here, DDM failed to extract full-length material from cell membranes. SMA offers an excellent alternative to canonical detergents, as these polymers directly extract membrane proteins and protein complexes from cell membranes inside a nanoparticle of annular lipids.[19, 21] Previous experiments in our group demonstrated that SMALPs were excellent vehicles for solubilization of Sm-PGTs.[23, 24] Recombinantly expressed Lg-PGTs from S. *enterica, T. thermophilus, E. coli*, and S. *pneumoniae* were readily solubilized in SMALP directly from isolated *E. coli* cell membranes. By utilizing the highly specific dual-Strep Tag for purification, PGT-containing SMALPs were purified in a single step. Purity and final yield of PGTs varied depending on the type of SMA polymer used for purification, with the combination of SMA30 and S. *enterica* WbaP demonstrating optimal yield and purity as judged by SDS-PAGE (Fig. 3). The advantage of SMALPs for purification of the Lg-PGTs described here is striking as shown by the comparison of SMA30 and DDM solubilization, side-by-side, on identical cell membrane extracts. None of the PGTs were stable or soluble in DDM micelles, while each PGT tested was successfully isolated in SMALP (Fig S1). Although all the proteins produced here displayed activity in isolated cell membranes, none of the Lg-PGTs were active in SMALP. SMALP-mediated influences on protein function are not unprecedented and may indicate the rigidity of the bilayer stabilized by the SMA prevents catalysis.[43] Alternatively, as PGTs require Mg^2+^ as a cofactor, the chelating nature of the maleic acid moiety found on SMA polymers may strip the Lg-PGTs of sufficient Mg^2+^ for catalysis.[25, 33, 34] Despite these limitations, SMALPs may still be used as a “shuttle” membrane mimetic to stably transfer Lg-PGTs from cell membrane to other systems such as proteoliposomes.[44]

Although S. *enterica* WbaP in SMA30 appeared highly pure by SDS-PAGE, the quality of the protein in SMALPs in solution remained unclear. As CryoEM optics, sample preparation techniques, and data processing strategies have continued to improve, characterization of small membrane proteins both in detergent or SMALP has become more feasible.[45, 46] A small screening dataset collected on a Talos Arctica microscope revealed S. *enterica* WbaP particles were readily distinguishable. Particles had a diameter ~120 Å, similar to that reported for SMALPs previously.[20] 2D classification of 189,401 particles extracted from 475 micrographs revealed classes with distinct features, including apparent C2 symmetry (Fig 5). The simplicity of the expression and purification protocol described here and the apparent high-quality of the protein produced are ideally suited for Cryo-EM of difficult membrane proteins. Future work will aim to resolve the structure of a Lg-PGT in SMALP to high resolution to better understand the molecular architecture of these enzymes in a native-like environment.

## CRediT authorship contribution statement

**Greg J. Dodge:** Conceptualized and carried out experiments, prepared samples and collected data for electron microscopy, analyzed and visualized data, generated figures, Writing – original draft, review & editing. **Hannah M. Bernstein:** Methodology, carried out experiments, analyzed and visualized data, Writing – review & editing. **Barbara Imperiali:** Conceptualization, Writing – review & editing, supervision, and funding acquisition.

## Data Availability

Any data, plasmids, or protocols will be made available upon request to authors.

### Funding

This work was funded by National Institutes of Health gran R01 GM131627 and GM039334 (to B.l), and F32 GM134576 (G.J.D.)

## Abbreviations

PGT: phosphoglycosyl transferase
SMALP: styrene maleic acid co-polymer liponanoparticle
CryoEM: cryogenic electron microscopy
LPS: lipopolysaccharide
CPS: capsular polysaccharide
EPS: exopolysaccharide
WTA: Wall teichoic acid
UndP: undecaprenol phosphate
Poly: polytopic
Mono: monotopic
TMH: transmembrane helix
RMH: re-entrant membrane helix
Sm: small
Bi: bifunctional
Lg: large
DUF: domain of unknown function
SSN: sequence similarity network
DDM: n-dodecyl-β-D-maltoside
LDAO: N.N-dimethyl-N-dodecylamine-N-oxide
DIBMA: di-isobutylene maleic acid
PSUP: pure solvent upper phase
IMAC: immobilized metal affinity chromatography
TEV: tobacco etch virus
SDS-PAGE: sodium dodecyl sulfate polyacrylamide gel electrophoresis

