## Supplementary Materials for "A generalizable protocol for expression and purification of membrane-bound bacterial phosphoglycosyl transferases in liponanoparticles"

^‡^Equal contributions

ORCID IDs

BI: 0000-0002-5749-7869

HMB: [0000-0003-0871-0376](https://orcid.org/0000-0003-0871-0376)

GJD: [0000-0002-6555-8350](https://orcid.org/0000-0002-6555-8350)

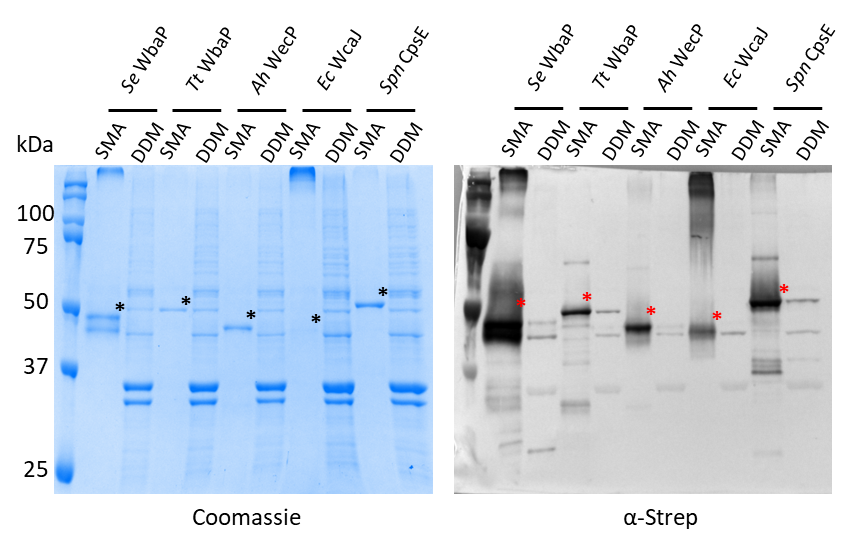

Figure S1 Comparison of SMA and DDM solubilization of selected Lg-PGTs. Left: SDS-PAGE gel with Coomassie staining. Right: Western blot using an α-strep 1° antibody. Asterisks indicate bands corresponding to full-length Lg-PGTs. Owing to the high isoelectric points, each PGT runs slightly below the anticipated molecular weight.

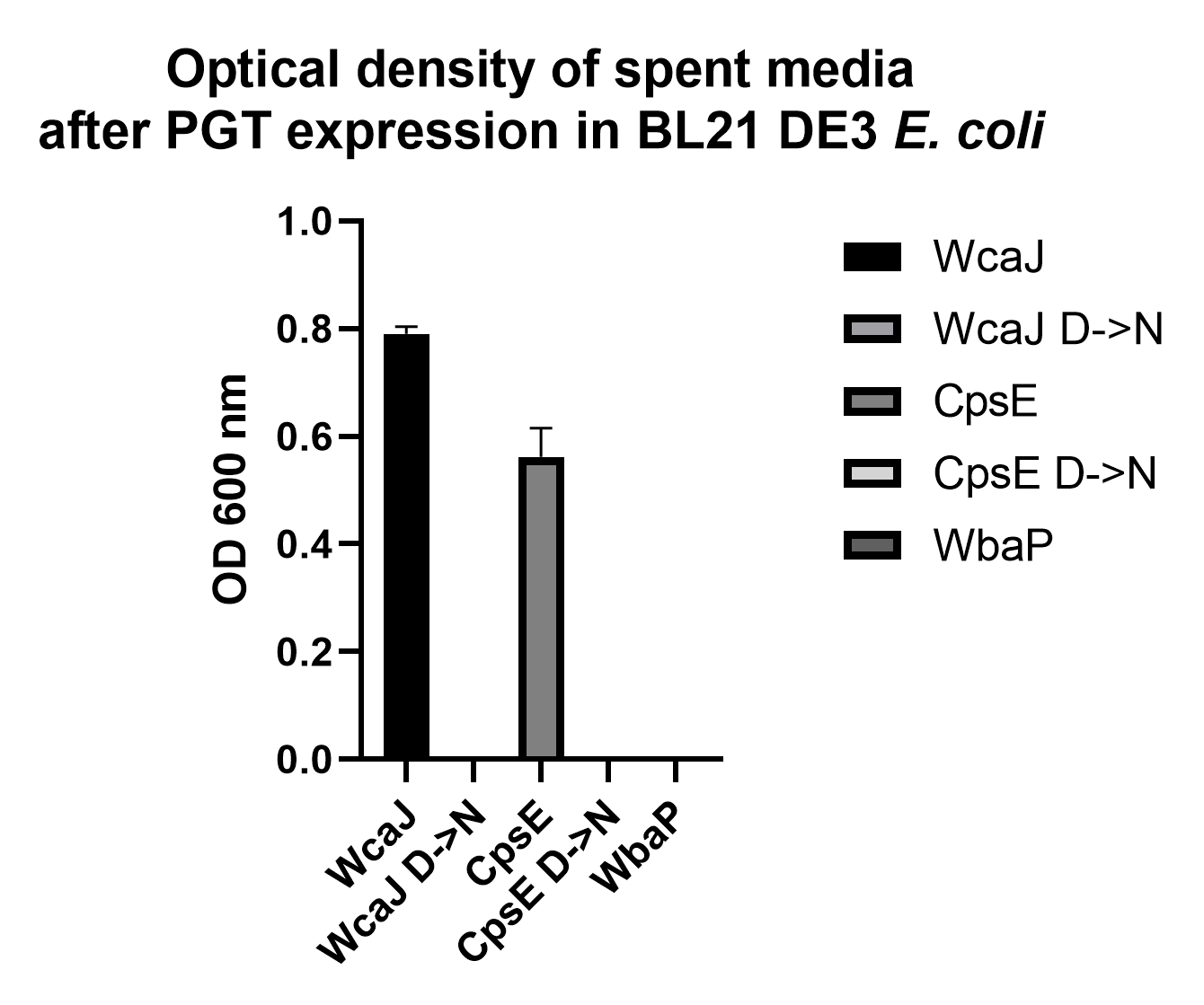

Figure S2 Analysis of spent media after expression of Lg-PGTs in BL21 DE3 E. coli. Both Wild-type Ec WcaJ and Spn CpsE caused spent media to have high turbidity. Media was also extremely viscous. This behavior was not observed when Se WbaP or D->N variants of WcaJ and CpsE were overexpressed.

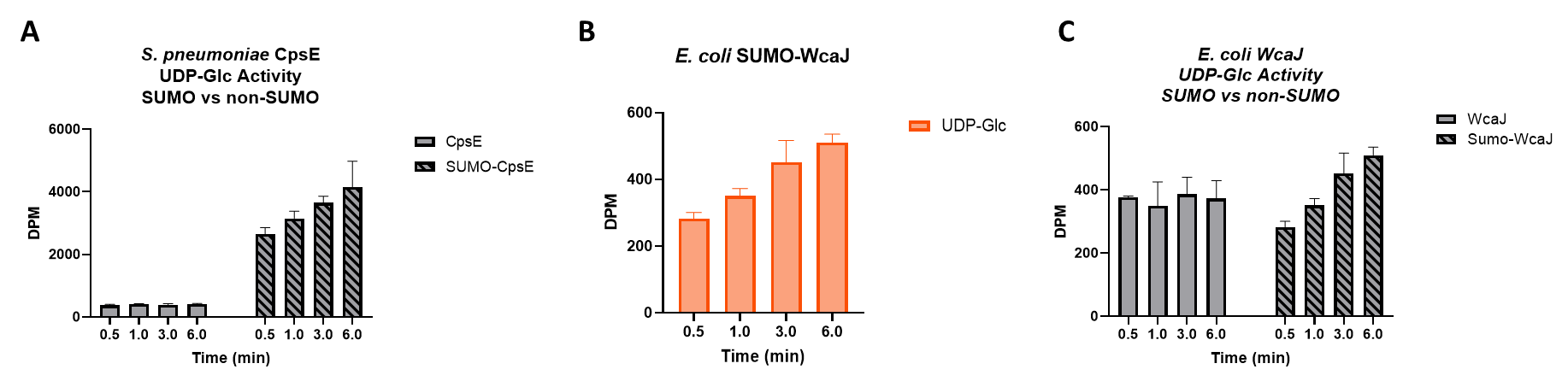

Figure S3 Activity of E. coli WcaJ in cell membranes utilizing UDP-Glc as substrate. A: Minimal UDP-Glc activity was observed for CpsE lacking a SUMO tag. B: A time-dependent increase in UDP-Glc transfer to UndP was observed for BL21 DE3 E. coli membranes containing SUMO-WcaJ. C: No increase in UDP-Glc transfer was observed over time for C43 E. coli membranes containing untagged WcaJ.

| [^3^H] Substrate | dpm | Specific Activity (Ci/mmol) |
| --- | --- | --- |
| UDP-Gal | 73,000 | 40 |
| UDP-GalNAc | 65,000 | 20 |
| UDP-Glc | 56,000 | 60 |
| UDP-Glc* | 200,000 | 6 |
| UDP-GlcNAc | 67,000 | 20 |

Table S1 [^3^H] UDP-sugar substrates used for Lg-PGT assays. *Used for characterization of SUMO-CpsE and SUMO-WcaJ.
